# Mnemonic brain state engagement is diminished in healthy aging

**DOI:** 10.1101/2024.08.12.607567

**Authors:** Isabelle L. Moore, Devyn E. Smith, Nicole M. Long

## Abstract

Healthy older adults typically show impaired episodic memory – memory for when and where an event oc-curred – but intact semantic memory – knowledge for general information and facts. As older adults also have difficulty inhibiting the retrieval of prior knowledge from memory, their selective decline in episodic memory may be due to a tendency to over engage the retrieval state, a brain state in which attention is focused internally in an attempt to access prior knowledge. The retrieval state trades off with the encoding state, a brain state which supports the formation of new memories. Therefore, episodic memory declines in older adults may be the result of differential engagement in mnemonic brain states. Our hypothesis is that older adults are biased toward a retrieval state. We recorded scalp electroencephalography while young, middle-aged and older adults performed a memory task in which they were explicitly directed to either encode the currently presented object stimulus or retrieve a previously presented, categorically-related object stimulus. We used multivariate pattern analysis of spectral activity to decode engagement in the retrieval vs. encoding state. We find that all three age groups can follow top-down instructions to selectively engage in encoding or retrieval and that we can decode mnemonic states for all age groups. However, we find that mnemonic brain state engagement is diminished for older adults relative to middle-aged adults. Our interpretation is that a combination of executive control deficits and a modest bias to retrieve modulates older adults’ mnemonic state engagement. Together, these findings suggest that differential mnemonic state engagement may underlie age-related memory changes.

## Introduction

In healthy aging, many aspects of cognition decline (Craik & Bialystok, 2006; Lindenberger, 2014), including memory (Rabin et al., 2015). Healthy older adults exhibit selective deficits in episodic memory – the memory for events within a spatiotemporal context (Tulving, 1972) – but typically have intact semantic memory – memory for facts and concepts (Wingfield & Kahana, 2002). For example, an older adult may be easily able to remember that *Water Lilies* was painted by Claude Monet, but may fail to remember the specific details of their trip to the Art Institute of Chicago during which they viewed the painting. This selective episodic memory deficit may be due to a tendency to engage a retrieval state, given older adults’ difficulty inhibiting the retrieval of prior knowledge from memory (Wynn, Ryan, & Moscovitch, 2020). In young adults, engaging the retrieval state (Tulving, 1983) trades off with engaging the encoding state (Long & Kuhl, 2019; Smith, Moore, & Long, 2022) and can impair later memory (Long & Kuhl, 2019). Age-related episodic memory deficits may therefore arise from differential mnemonic state engagement. The aim of the present study is to investigate the extent to which older adults engage encoding and retrieval states.

Relative to young adults, older adults have impaired episodic memory and intact semantic memory. Older adults have difficulty forming and accessing episodic associations – connections between events that occur close in time or space (Wilkniss, Jones, Korol, Gold, & Manning, 1997; Golomb, Peelle, Addis, Kahana, & Wingfield, 2008), but typically have no such difficulty with semantic associations (Amer, Giovanello, Nichol, Hasher, & Grady, 2019) – connections between events that share meaning. For instance, when studying realistic and unrealistic prices for grocery store items, older adults show memory comparable to young adults only for those items that are priced realistically, suggesting that older adults rely on prior knowledge to support memory judgments (Amer, Giovanello, Grady, & Hasher, 2018). Older adults also rely on semantic memory rather than episodic memory to guide decision-making, even when episodic memory is more task-relevant (Lalla, Tarder-Stoll, Hasher, & Duncan, 2022). Together, these findings suggest that older adults may over rely on semantic information.

An overreliance on retrieving semantic information may prevent the encoding of episodic information. In young adults, attending to semantic associations can negatively impact the formation of episodic associations (Long & Kahana, 2017). Memory for semantic vs. episodic details – e.g. “Claude Monet was a French Impressionist” vs. “We visited the Art Institute of Chicago last Tuesday at noon” – are negatively associated, an effect which increases with age (Devitt, Addis, & Schacter, 2017). There is an overall shift toward semantic memory across the lifespan (Ofen & Shing, 2013) along with increases in default mode network (DMN) connectivity (Staffaroni et al., 2018), a brain network known to support semantic processing (Binder, Desai, Graves, & Conant, 2009) among other internally-directed processes (Buckner & DiNicola, 2019). Given that in young adults DMN activation impairs episodic memory formation (Kim, 2011), older adults’ tendency to engage in DMN-mediated semantic retrieval may prevent or diminish episodic encoding.

Older adults may engage in retrieval at the expense of encoding. Older adults are biased toward pattern completion – reactivation of a memory trace via the hippocampus from incomplete sensory input (McClelland, McNaughton, & O’Reilly, 1995) – whereby older adults are more likely than young adults to identify novel stimuli with low similarity to studied items as familiar (Vieweg, Stangl, Howard, & Wolbers, 2015). When older adults retrieve prior information to make predictions about future events, the retrieved information may be strengthened at the expense of encoding new events (Stawarczyk, Wahlheim, & Zacks, 2023). Memory encoding and memory retrieval constitute neurally distinct brain states (Hasselmo, Bodelon, & Wyble, 2002; Hasselmo, 2005) – spatially distributed and temporally sustained neural activity patterns (Beaty et al., 2018; Kay & Frank, 2019; Tang, Rothbart, & Posner, 2012) – which trade off such that they cannot be engaged in simultaneously. Encoding and retrieval states can be distinguished through neural activity patterns recorded via scalp electroencephalography (Long & Kuhl, 2019; Smith et al., 2022; Hong, Moore, Smith, & Long, 2023; Smith & Long, 2024) and functional magnetic resonance imaging (Long & Kuhl, 2021). An investigation of microstates, global patterns of voltage scalp topography that are stable over successive short time periods (Michel & Koenig, 2018), revealed that sustained engagement of microstate E – previously linked to the DMN (Custo et al., 2017; Bréchet et al., 2019) – characterizes the retrieval state (Hong et al., 2023). Young adults engage the retrieval state in response to top-down demands to retrieve (Smith & Long, 2024), but also automatically when experiences overlap in time (Smith et al., 2022), suggesting that the retrieval state may be engaged even when counter to top-down goals. As older adults have a diminished ability to inhibit retrieval of semantic information (Wynn et al., 2020), they may be particularly susceptible to engaging the retrieval state, which in turn will impair encoding. Thus, an older adult may fail to remember their specific trip to the Art Institute because when they encountered *Water Lilies*, they automatically retrieved their prior knowledge about Claude Monet (e.g. Impressionism, France, gardening) which prevented them from encoding the current experience.

Our hypothesis is that older adults are biased toward the retrieval state. To test our hypothesis, we conducted a scalp electroencephalography study in which young (18-35), middle-aged (36-59) and older adult (60-85) participants studied two lists of object images. The second list was comprised of images that were semantically associated with images from the first list (i.e., each image in a “pair” was drawn from the same object category). During List 2, participants were explicitly instructed to either encode the currently presented object image or retrieve the categorically related object image from List 1. Following study, participants completed a two-alternative forced-choice recognition test of all images from both lists. We used multivariate pattern analyses to assess engagement in memory encoding and memory retrieval brain states. To the extent that older adults are biased to retrieve, we should find differential mnemonic brain state engagement for older adults compared to young and/or middle-aged adults.

## Materials and Methods

### Participants

Forty young adult (34 female, age range = 18-37, mean age = 20.3 years), forty middle-aged adult (34 female, age range = 39-59, mean age = 51.4 years) and forty older adult (26 female, age range = 60-85, mean age = 70.5 years) fluent English speakers from the University of Virginia (UVA) community participated. Young adult participants (YAs) were recruited from flyers posted around the UVA campus and surrounding areas. Middle-aged adults (MAs) and older adults (OAs) were recruited through the Virginia Cognitive Aging Project, a longitudinal study of cognitive aging in adults across a wide age range (Salthouse, 2011). The YA dataset has been previously reported (Smith et al., 2022); albeit with a slight modification as we have removed an additional participant (see below). Our enrollment targets for the MA and OA datasets were based on this YA dataset and are described in the pre-registration report of this study (https://osf.io/8ezpr/). All participants had normal or corrected-to-normal vision. Informed consent was obtained in accordance with the UVA Institutional Review Boards for Social and Behavioral Research and Health Sciences Research, and participants were compensated for their participation. All MA and OA participants completed the Montreal Cognitive Assessment (MoCA); any participant who scored less than 26 was excluded, based on previous literature (Nasreddine et al., 2005).

Four YA participants were excluded from the final dataset: three who had been excluded in our previous report (Smith et al., 2022) for poor task performance, prior knowledge of the task, or poor signal quality, and one additional participant who was older than our YA upper age cut off of 35 at the time of testing. Four MA participants were excluded from the final dataset: one who had poor task performance (recognition accuracy *<* 3 SDs of the mean of the full MA dataset), one because of permanent hair extensions that precluded scalp access, resulting in poor signal quality (impedances were above the threshold of 50 kΩ), and two who scored lower than 26 on the MoCA. Two OA participants were excluded from the final dataset: one who had poor task performance (recognition accuracy *<* 3 SDs of the mean of the full OA dataset) and one who scored lower than 26 on the MoCA. Thus, data are reported for the remaining 36 YA, 36 MA and 38 OA participants. The raw, de-identified data and the associated experimental and analysis codes used in this study will be made available for access via the Long Term Memory laboratory website upon publication of this manuscript (https://longtermmemorylab.com).

## Materials

### Mnemonic State Task Experimental Design

Stimuli consisted of 576 object pictures, drawn from an image database with multiple exemplars per object category (Konkle, Brady, Alvarez, & Oliva, 2010). From this database, we chose 144 unique object categories and 4 exemplars from each category. For each participant, one exemplar in a set of four served as a List 1 object, one as a List 2 object, and the two remaining exemplars served as lures for the recognition phase. Object condition assignment was randomly generated for each participant.

#### General Overview

In each of eight runs, participants viewed two lists containing 18 object images. For the first list, each object was new (List 1 objects). For the second list (List 2 objects), each object was again new, but was categorically related to an object from the first list. For example, if List 1 contained an image of an apple, List 2 would contain an image of a different apple (Figure 1). During List 1, participants were instructed to encode each new object. During List 2, however, each trial contained an instruction to either encode the current object (e.g., the new apple) or to retrieve the corresponding object from List 1 (the old apple). Following eight runs, participants completed a two-alternative forced-choice recognition test that separately assessed memory for List 1 and List 2 objects.

**Figure 1.**
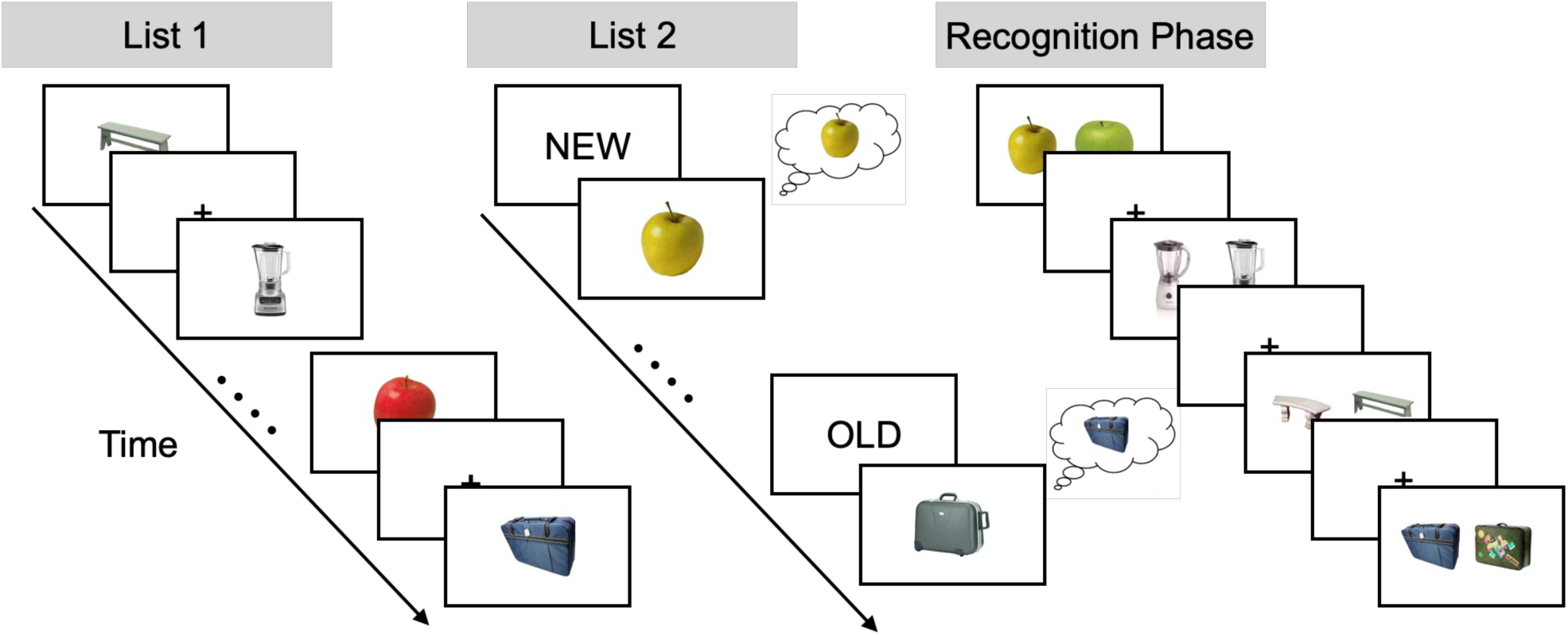
Task design. During List 1, participants studied individual objects (e.g., apple, suitcase). During List 2, participants saw novel objects that were from the same categories as the objects shown in List 1 (e.g., a new apple, a new suitcase). Preceding each List 2 object was an OLD instruction cue or NEW instruction cue. The OLD cue signaled that participants were to retrieve the corresponding object from List 1 (e.g., the old suitcase). The NEW cue signaled that participants were to encode the current object (e.g., the new apple). Each run of the experiment contained a List 1 and List 2; object categories (e.g., apple) were not repeated across runs. After eight runs, participants completed a two-alternative forced-choice recognition test that tested memory for each List 1 and List 2 object. On each trial, a previously presented object, either from List 1 or List 2, was shown alongside a novel lure from the same category. The participant’s task was to choose the previously presented object. List 1 and List 2 objects were never presented together.

#### List 1

On each trial, participants saw a single object presented for 2000 ms followed by a 1000 ms inter-stimulus interval (ISI). Participants were instructed to study the presented object in anticipation of a later memory test.

#### List 2

On each trial, participants saw a cue word, either “OLD” or “NEW” for 2000 ms. The cue was followed by presentation of an object for 2000 ms, which was followed by a 1000 ms ISI. All objects in List 2 were non-identical exemplars drawn from the same category as the objects presented in the immediately preceding List 1. That is, if a participant saw a suitcase and an apple during List 1, a different suitcase and a different apple would be presented during List 2. On trials with a “NEW” instruction (encode trials), participants were to encode the presented object. On trials with an “OLD” instruction (retrieve trials), participants tried to retrieve the categorically related object from the preceding List 1. Importantly, this design prevented participants from completely ignoring List 2 objects following “OLD” instructions in that they could only identify the to-be-retrieved object category by processing the List 2 object.

Participants completed eight runs with two lists in each run (List 1, List 2). Participants viewed 18 objects per list, yielding a total of 288 object stimuli from 144 unique object categories. Participants did not make a behavioral response during either List 1 or 2. Following the eight runs, participants completed a two-alternative forced choice recognition test.

#### Recognition Phase

Following the eight runs, participants completed the recognition phase. On each trial, participants saw two exemplars from the same object category (e.g. two apples; Figure 1). One object had previously been encountered either during List 1 or 2. The other object was a lure and had not been presented during the experiment. Because both test probes were from the same object category, participants could not rely on familiarity or gist-level information to make their response (Brainerd & Reyna, 2002). Trials were self-paced and participants selected (via button press) the previously presented object. Trials were separated by a 1000 ms ISI. There were a total of 288 recognition trials (corresponding to the 288 total List 1 and 2 objects presented in the experiment). Note: List 1 and List 2 objects never appeared in the same trial together, thus participants never had to choose between two previously presented objects. List 1 and List 2 objects were presented randomly throughout the test phase.

### EEG Data Acquisition and Preprocessing

Electroencephalography (EEG) recordings were collected using a BrainVision system and an ActiCap equipped with 64 Ag/AgCl active electrodes positioned according to the extended 10-20 system. All electrodes were digitized at a sampling rate of 1000 Hz and were referenced to electrode FCz. Offline, electrodes were later converted to an average reference. Impedances of all electrodes were kept below 50 kΩ. Electrodes that demonstrated high impedance or poor contact with the scalp were excluded from the average reference calculations; however, all electrodes were included in all subsequent analysis steps following re-referencing. Bad electrodes were determined by voltage thresholding (see below).

Custom Python codes were used to process the EEG data. We applied a high pass filter at 0.1 Hz, followed by a notch filter at 60 Hz and harmonics of 60 Hz to each participant’s raw EEG data. We then performed three preprocessing steps (Nolan, Whelan, & Reilly, 2010) to identify electrodes with severe artifacts. First, we calculated the mean correlation between each electrode and all other electrodes as electrodes should be moderately correlated with other electrodes due to volume conduction. We *z*-scored these means across electrodes and rejected electrodes with *z*-scores less than -3. Second, we calculated the variance for each electrode as electrodes with very high or low variance across a session are likely dominated by noise or have poor contact with the scalp. We then *z*-scored variance across electrodes and rejected electrodes with a *|z| >* = 3. Finally, we expect many electrical signals to be autocorrelated, but signals generated by the brain versus noise are likely to have different forms of autocorrelation. Therefore, we calculated the Hurst exponent, a measure of long-range autocorrelation, for each electrode and rejected electrodes with a *|z| >* = 3. Electrodes marked as bad by this procedure were excluded from the average re-reference. We then calculated the average voltage across all remaining electrodes at each time sample and re-referenced the data by subtracting the average voltage from the filtered EEG data. We used wavelet-enhanced independent component analysis (Castellanos & Makarov, 2006) to remove artifacts from eyeblinks and saccades.

### EEG Data Analysis

We applied the Morlet wavelet transform (wave number 6) to the entire EEG time series across electrodes, for each of 46 logarithmically spaced frequencies (2-100 Hz; Smith et al., 2022). After log-transforming the power, we downsampled the data by taking a moving average across 100 ms time intervals from 4000 ms preceding to 4000 ms following object presentation during List 1 and List 2. We slid the window every 25 ms, resulting in 317 time intervals (80 non-overlapping). Power values were then *z*-transformed by subtracting the mean and dividing by the standard deviation power. Mean and standard deviation power were calculated across all List 1 and List 2 objects across time points for each frequency.

### Pattern Classification Analyses

Pattern classification analyses were performed using penalized (L2) logistic regression (penalty parameter = 1), implemented via the sklearn linear model module in Python. Classifier performance was assessed in two ways. “Classification accuracy” represented a binary coding of whether the classifier successfully guessed the instruction condition. We used classification accuracy for general assessment of classifier performance (i.e., whether encode/retrieve instructions could be decoded). “Classifier evidence” was a continuous value reflecting the logit-transformed probability that the classifier assigned the correct instruction for each trial. Classifier evidence was used as a trial-specific, continuous measure of mnemonic state information, which we used to assess the degree to which retrieval state evidence varied across time as a function of the task instruction (encode, retrieve) and age group. Positive classifier evidence values indicate more evidence for a retrieval state and negative classifier evidence values indicate more evidence for an encoding state.

We trained within-participant classifiers to discriminate List 2 encode vs. retrieve trials based on a feature space comprised of all 63 electrodes *×* 46 logarithmically spaced frequencies ranging from 2 to 100 Hz. Before pattern classification analyses were performed, we performed an additional round of *z*-scoring across features (electrodes and frequencies) to eliminate trial-level differences in spectral power (Smith et al., 2022). For each participant, we used leave-one-run-out cross-validated classification in which the classifier was trained to discriminate encode from retrieve instructions for seven of the eight runs and tested on the held-out run. For classification analyses in which we assessed classifier accuracy, we averaged z-transformed power values over the 2000 ms stimulus interval. We conducted four separate classification analyses on z-transformed power averaged across four 500 ms intervals (0-500, 500-1000, 1000-1500, 1500-2000) spanning the stimulus interval. We evaluated retrieval evidence as a function of instruction (encode, retrieve) across and within each of these intervals.

### Cross Study Memory State Decoding

We used an independently-validated mnemonic state classifier to test memory state engagement across MA and OA groups with a single training set, as within-participant classification can be driven by different features for different participants. We conducted three stages of classification using the same methods as in our prior work (Long, 2023; Smith & Long, 2024; Wheelock & Long, 2024). First, we conducted within participant leave-one-run-out cross-validated classification (penalty parameter = 1) on a set of participants who completed the mnemonic state task which includes the YA participants reported here (N = 103, see ref. Hong et al., 2023 for details). The classifier was trained to distinguish encoding vs. retrieval states based on spectral power averaged across the 2000 ms stimulus interval during List 2 trials. For each participant, we generated true and null classification accuracy values. We permuted condition labels (encode, retrieve) for 1000 iterations to generate a null distribution for each participant. Any participant whose true classification accuracy fell above the 90th percentile of their respective null distribution was selected for further analysis (N = 37). Second, we conducted leave-one-participant-out cross-validated classification (penalty parameter = 0.0001) on the selected participants to validate the mnemonic state classifier and obtained classification accuracy of 60% which is significantly above chance (*t* _36_ = 6.000, *p <* 0.0001), indicating that the cross-participant mnemonic state classifier is able to distinguish encoding and retrieval states. Finally, we applied the cross-participant mnemonic state classifier to the MA and OA List 2 data, specifically in 100 ms intervals across the 2000 ms stimulus interval. Because the cross-participant mnemonic state classifier was trained on data partially comprised of the YA data reported here, we did not apply it to the YA data. We extracted classifier evidence, the logit-transformed probability that the classifier assigned a given trial a label of encoding or retrieval. This approach provides a trial-level estimate of memory state evidence that is not constrained by participant-level differences in feature weights, enabling us to more directly compare mnemonic state engagement across the MA and OA groups.

### Statistical Analyses

We used mixed effects ANOVAs and paired or independent samples *t* -tests to assess the effect of encode/retrieve instruction on behavioral memory performance. We used a 3 *×* 2 *×* 2 ANOVA with age (YA, MA, OA), instruction (encode, retrieve), and list (first, second) as factors.

To evaluate classification accuracy for MAs and OAs, we used paired-sample *t* -tests to compare classification accuracy across participants within each group to chance decoding accuracy, as determined by permutation procedures. Namely, for each participant, we shuffled the condition labels of interest (e.g., “encode” and “retrieve” for the instruction classifier) and then calculated classification accuracy. We repeated this procedure 1000 times for each participant and then averaged the 1000 shuffled accuracy values for each participant. These mean values were used as participant-specific empirically derived measures of chance accuracy. We used paired samples *t* -tests to compare the observed (unshuffled) accuracy values to the shuffled accuracy values.

To assess whether classification accuracy varies across age group, we used a 1 *×* 3 ANOVA with age (YA, MA, OA) as a factor. The dependent variable was classification accuracy from the classifier trained and tested on spectral data averaged over the 2000 ms stimulus interval.

To assess the time course and strength of classifier evidence as a function of age, we used mixed effects ANOVAs. We used a 3 *×* 2 *×* 4 ANOVA with age (YA, MA, OA), instruction (encode, retrieve), and time interval (four 500 ms intervals spanning the stimulus interval) as factors.

## Results

### Middle-aged and older adults selectively encode and retrieve in response to top-down instructions

We first sought to determine the extent to which middle-aged (MAs) and older adults (OAs) can follow task instructions to engage in encoding and retrieval in comparison to young adults (YAs; Long & Kuhl, 2019; Smith et al., 2022). Specifically, we tested whether instructions influenced recognition task performance across age groups (Figure 2). Following our pre-registration, we conducted a 3*×*2*×*2 mixed effects ANOVA to evaluate the effects of age group (YAs, MAs, OAs), instruction type (encode, retrieve) and list (first, second) on recognition accuracy. A two-way interaction between list and instruction type, with no three-way interaction between list, instruction type, and age group, would indicate that instruction type has a similar impact on memory behavior for all three age groups. We report the full ANOVA results in Table 1 and highlight key findings here. We find a significant two-way interaction between list and instruction type, driven by greater recognition accuracy for List 2 objects with an encode instruction (*M* = 0.84, *SD* = 0.09) compared to List 2 objects with a retrieve instruction (*M* = 0.80, *SD* = 0.08; *t* _109_ = 4.91, *p <* 0.001, Cohen’s *d* = 0.42). The three-way interaction of list, instruction type and age group was not significant and Bayes Factor analysis revealed that a model without the three-way interaction term (H_0_) is preferred to a model with the three-way interaction term (H_1_; BF_10_ = 0.07, strong evidence for H_0_). These results suggest that like YAs, MAs and OAs can follow instruction cues to selectively encode or retrieve.

**Figure 2.**
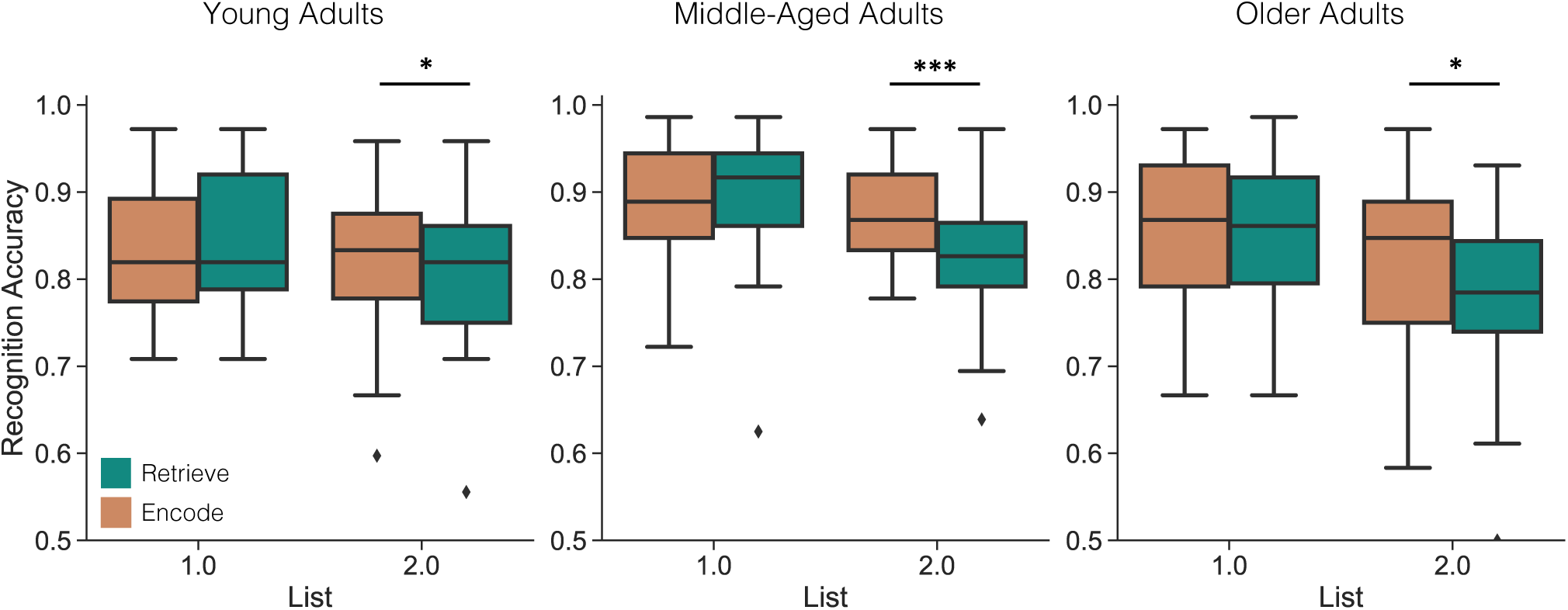
Influence of age and mnemonic instructions on memory behavior. We assessed recognition accuracy as a function of list (1, 2) and instruction (orange represents encode; teal represents retrieve) for each age group. The boxes denote interquartile range, the horizontal bars denote median, and the whiskers denote minimum and maximum values. We find a significant interaction between list and instruction (*p <* 0.001) driven by greater accuracy for List 2 objects presented with an encode compared with a retrieve instruction. The three-way interaction of list, instruction type and age group was not significant (*p* = 0.399). \**p <* 0.05, \*\*\**p <* 0.001.

**Table 1.** Analysis of variance for the effect of instruction type, age group, and list on recognition accuracy.

| Effect | df | *F* | *p* | $\eta_p^2$ |
| --- | --- | --- | --- | --- |
| Main effect of age group | (2,107) | 5.88 | **0.004** | 0.010 |
| Main effect of list | (1,107) | 34.09 | **<0.001** | 0.24 |
| Main effect of instruction type | (1,107) | 14.60 | **<0.001** | 0.12 |
| Interaction of age group × list | (2,107) | 2.27 | 0.108 | 0.04 |
| Interaction of age group × instruction type | (2,107) | 1.08 | 0.342 | 0.02 |
| Interaction of instruction type × list | (1,107) | 18.87 | **<0.001** | 0.15 |
| Interaction of age group × instruction type × list | (2,107) | 0.93 | 0.399 | 0.02 |

*Bold values indicate $p < 0.05$ .*

### Middle-aged and older adults differentially engage mnemonic states

Having shown behavioral evidence that MAs and OAs can follow instructions to selectively engage in encoding and retrieval, we next sought to determine the extent to which they engage mnemonic brain states in response to the instruction cues. Following our pre-registration, we conducted a multivariate pattern classification analysis in which we trained within-participant, leave-one-run-out classifiers to discriminate encode versus retrieve List 2 trials based on a feature space comprised of 63 electrodes and 46 frequencies ranging from 2-100 Hz averaged over the 2000 ms stimulus interval (Figure 3). For all age groups, mean classification accuracy (YAs: *M* = 54.09%; MAs: *M* = 58.24%; OAs: *M* = 54.81%) was significantly greater than chance (as determined through a permutation procedure; all *p*s *<* 0.01). These results suggest that like YAs, both MAs and OAs can shift between encoding and retrieval states in a goal-directed manner.

**Figure 3.**
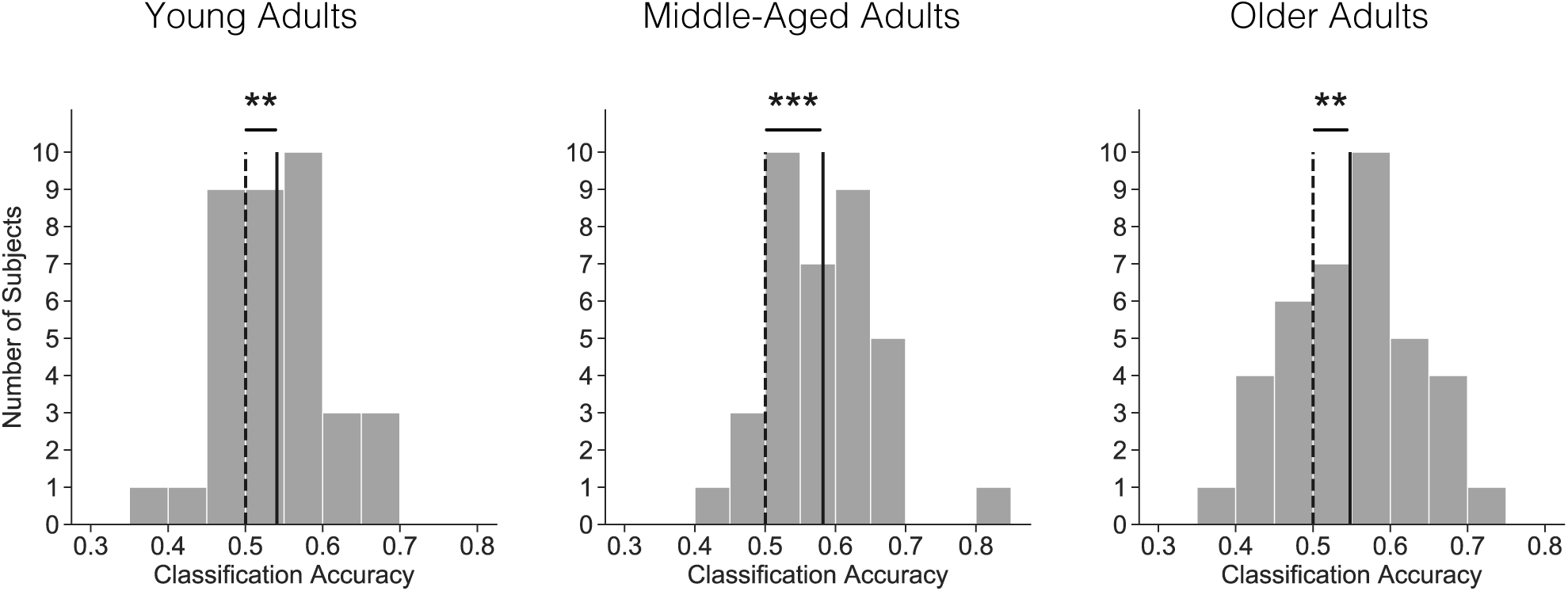
Mnemonic state classification accuracy by age group. We conducted L2-logistic regression classification to discriminate encode versus retrieve trials during List 2 separately for each age group. Within-participant classifiers were trained and tested on a feature space of 63 electrodes and 46 frequencies averaged across the 2000 ms stimulus interval. Mean classification accuracy across all participants within each age group (solid vertical line) is shown along with a histogram of classification accuracies for individual participants (gray bars) and mean classification accuracy for permuted data across all participants (dashed vertical line). Mean classification accuracy was 54.09% for young adults, 58.24% for middle-aged adults, and 54.81% for older adults, each of which differed significantly from chance (all *p*s *<* 0.01). \*\**p <* 0.01, \*\*\**p <* 0.001.

Our next goal was to determine if classification accuracy varies as a function of age. Following our pre-registration, we conducted a 1*×*3 mixed effects ANOVA to evaluate the effect of age group (YAs, MAs, OAs) on classification accuracy. The main effect of age group did not reach significance (*F* _2,107_ = 2.83, *p* = 0.064, *η_p_*^2^ = 0.05) and Bayes factor analysis yielded a value of 0.84, indicating anecdotal evidence for the null hypothesis that there is no difference in classification accuracy on the basis of age group. However, given the weak Bayes factor value and the possibility that group differences are present even when an ANOVA does not yield differences (Midway, Robertson, Flinn, & Kaller, 2020), we opted to perform post-hoc *t* -tests. We found that classification accuracy is significantly greater for MAs (*M* = 58.24%, *SD* = 7.80%) compared to YAs (*M* = 54.09%, *SD* = 7.20%; *t* _70_ = 2.31, *p* = 0.024, Cohen’s *d* = 0.55) and numerically greater for MAs compared to OAs (*M* = 54.81%, *SD* = 8.37%; *t* _72_ = 1.80, *p* = 0.077, Cohen’s *d* = 0.42), with no difference in classification accuracy between YAs and OAs (*t* _72_ = 0.39, *p* = 0.699, Cohen’s *d* = 0.09). These results indicate that we can successfully distinguish encode from retrieve trials in MAs and OAs and that YAs and OAs show comparable decoding performance.

Although we hypothesized that OAs are biased toward a retrieval state, the similar decoding performance across YAs and OAs – as well as our behavioral findings – suggests that OAs do engage the encoding state when instructed to encode. However, classification accuracy may obscure both moment-to-moment differences in memory state engagement between age groups and overall differences in memory state engagement. Therefore, we next sought to assess classifier evidence across the 2000 ms stimulus interval. Classifier evidence provides a continuous measure of the degree to which brain patterns resemble either a retrieval state (positive evidence) or an encoding state (negative evidence). Thus, we can use classifier evidence to measure the extent to which OAs compared to YAs and MAs engage in mnemonic states and whether this engagement fluctuates over time. We have previously found that YAs show ro-bust dissociations in retrieval state evidence across encode and retrieve trials late in the stimulus interval (around 1000 ms following stimulus onset; Smith et al., 2022).

Based on our prior work and following our pre-registration, we assessed classifier evidence across four 500 ms time intervals spanning the stimulus interval (Figure 4A). We conducted a 3*×*2*×*4 mixed effects ANOVA to evaluate the effects of age group (YA, MA, OA), instruction type (encode, retrieve) and time interval (0-500, 500-1000, 1000-1500, 1500-2000 ms) on classifier evidence. A three-way interaction between age group, instruction type, and time interval would indicate differential memory state engagement by age that fluctuates over time. Alternatively, a two-way interaction between age group and instruction type, with no three-way interaction, would indicate that age differences in memory state engagement are consistent throughout the stimulus interval. We report the results of this ANOVA in Table 2 and highlight the key findings here. We find a significant interaction between age group and instruction type. The three-way interaction between age, instruction, and time interval was not significant. Bayes Factor analysis revealed that a model without the three-way interaction term (H_0_) is preferred to a model with the three-way interaction term (H_1_; BF_10_ = 0.08, strong evidence for H_0_).

**Figure 4.**
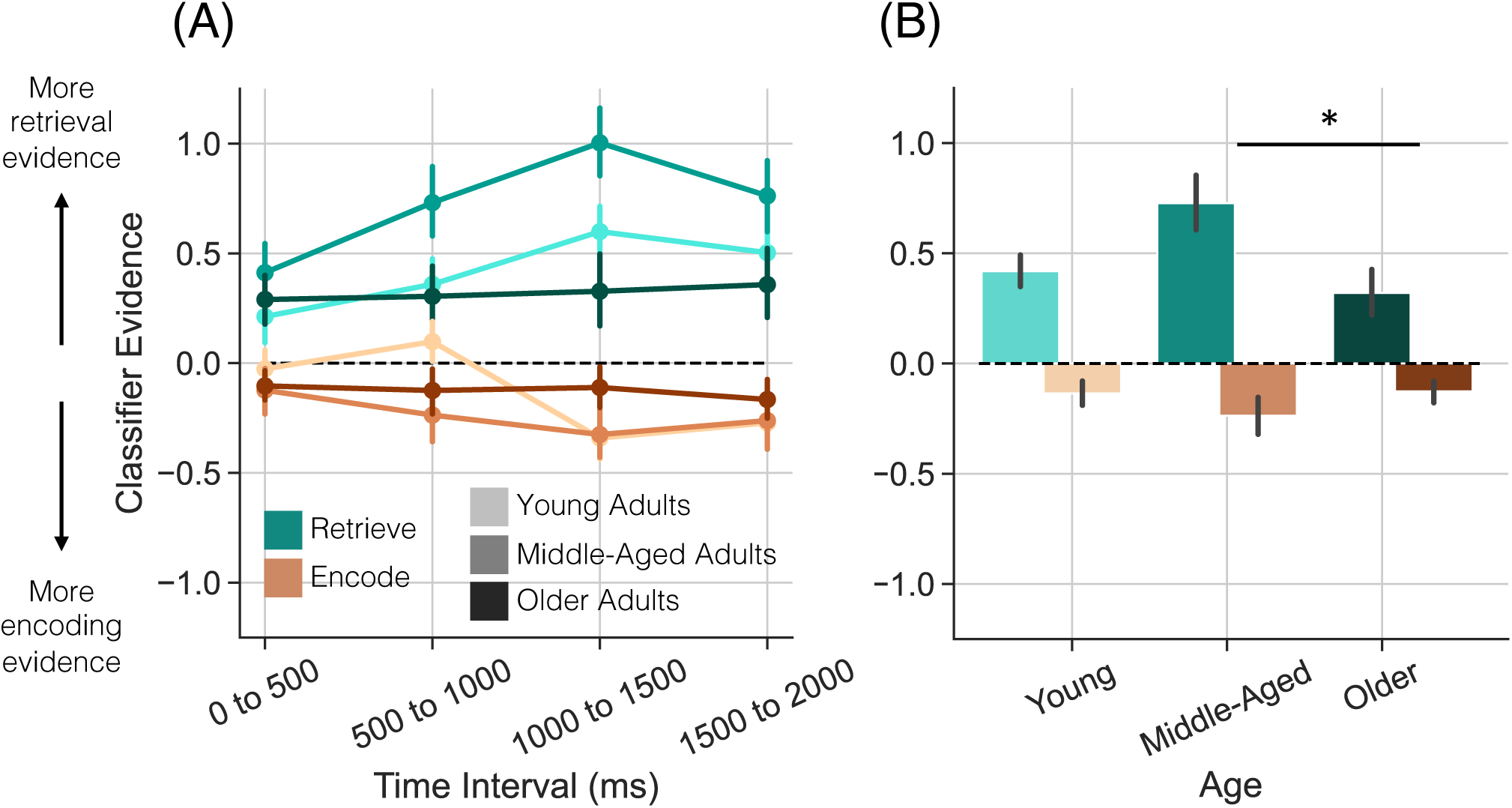
Classifier evidence by age group. We trained and tested four sets of within-participant classifiers on four 500 ms time intervals within the 2000 ms stimulus interval separately for each age group. We assessed classifier evidence as a function of age group (young, lightest; middle-aged, intermediate; older, darkest), and instruction type (encode, orange; retrieve, teal). Positive values indicate greater evidence for retrieval; negative values indicate greater evidence for encoding. Error bars denote standard error of the mean. **(A)** Time interval (four 500 ms intervals) is included as a factor. We find a significant interaction between age group and instruction type (*p* = 0.032). **(B)** Classifier evidence averaged over time. The difference between retrieve and encode trials is significantly greater for middle-aged compared to older adults (*p* = 0.026). \**p <* 0.05

**Table 2.**
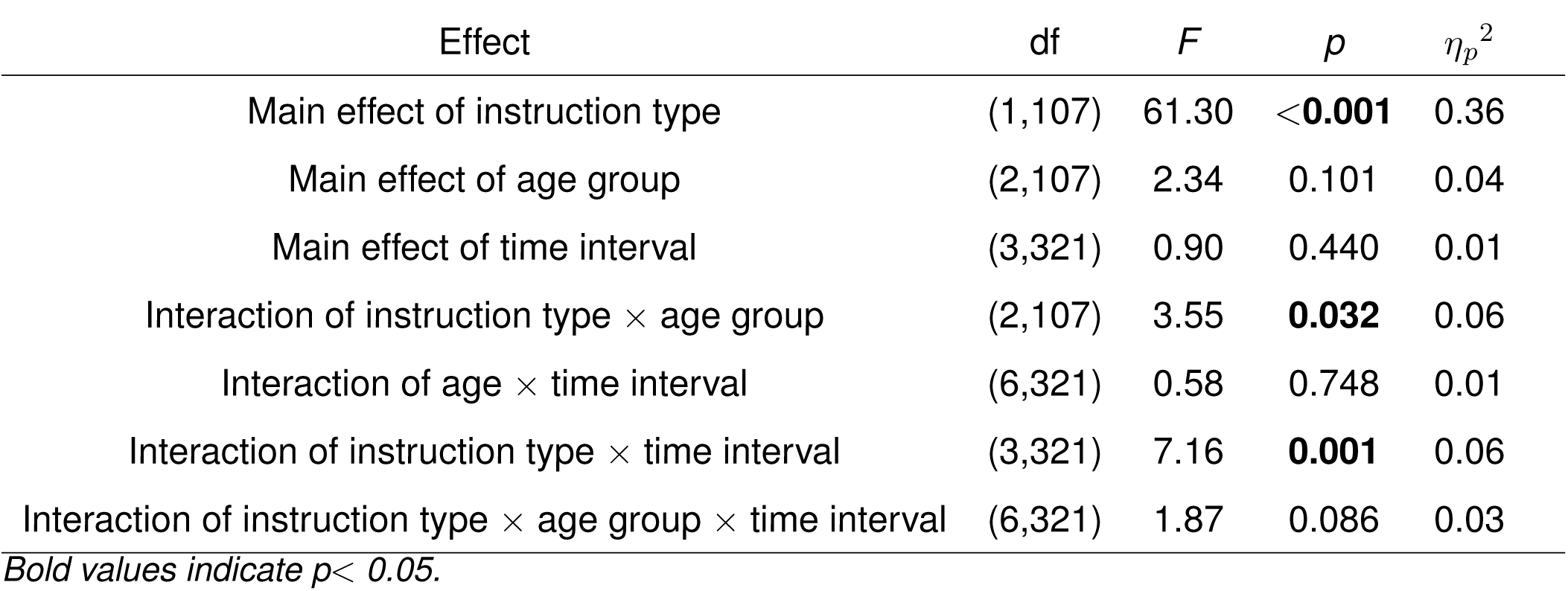
Analysis of variance for the effect of instruction type, age group, and time interval (500 ms) on retrieval evidence.

| Effect | df | *F* | *p* | $\eta_p^2$ |
| --- | --- | --- | --- | --- |
| Main effect of instruction type | (1,107) | 61.30 | **<0.001** | 0.36 |
| Main effect of age group | (2,107) | 2.34 | 0.101 | 0.04 |
| Main effect of time interval | (3,321) | 0.90 | 0.440 | 0.01 |
| Interaction of instruction type × age group | (2,107) | 3.55 | **0.032** | 0.06 |
| Interaction of age × time interval | (6,321) | 0.58 | 0.748 | 0.01 |
| Interaction of instruction type × time interval | (3,321) | 7.16 | **0.001** | 0.06 |
| Interaction of instruction type × age group × time interval | (6,321) | 1.87 | 0.086 | 0.03 |

*Bold values indicate $p < 0.05$ .*

Given the significant interaction between instruction type and age, we next directly compared average classifier evidence *differences* across age groups (Figure 4B). We calculated evidence difference scores (retrieve instruction trials -encode instruction trials) for each age group and performed three two-sample *t* -tests comparing each set of age groups. Positive values indicate more retrieval evidence for retrieve compared to encode trials. Difference scores were significantly lower for OAs (*M* = 0.45, *SD* = 0.82) compared to MAs (*M* = 0.96, *SD* = 1.10; *t* _72_ = 2.27, *p* = 0.026, Cohen’s *d* = 0.53). YA difference scores (*M* = 0.55, *SD* = 0.60) did not significantly differ from either MAs (*t* _70_ = 1.94, *p* = 0.057, Cohen’s *d* = 0.46) or OAs (*t* _72_ = 0.63, *p* = 0.529, Cohen’s *d* = 0.15). Together, these results suggest that although OAs can selectively engage mnemonic states, they do so to a lesser degree as compared to MAs.

### Older adults show diminished mnemonic brain state engagement relative to middle-aged adults

Our results suggest that although OAs can selectively engage memory encoding and memory retrieval states, the difference in engagement between the two mnemonic states is diminished relative to MAs. However, within-participant classification – although widely used – presents several challenges when attempting to interpret group-level differences. First, there is the possibility that the diminished disso-ciations in memory state evidence for OAs is due to issues with signal and/or data quality, rather than actual age-driven cognitive differences. Second, OAs may recruit different mechanisms when engaging in mnemonic states, as the exact features leveraged by each within-participant classifier can vary on a participant-by-participant basis. That is, although we find above chance classification for each age group, the features underlying classification success are not necessarily the same across age groups. Indeed, prior EEG work has shown that spectral signals change across the lifespan (Sander, Werkle-Bergner, & Lindenberger, 2012; Voytek et al., 2015; Tran, Hoffner, LaHue, Tseng, & Voytek, 2016; Healey & Kahana, 2020; Tran, Rolle, Gazzaley, & Voytek, 2020). In order to circumvent these limitations, we next utilized an independent dataset to assess mnemonic state engagement in MAs and OAs.

We conducted an exploratory analysis in which we applied an independently-validated mnemonic state classifier to the current MA and OA data. This classifier was trained to distinguish encoding and retrieval states from an independent group of participants who completed the mnemonic state task. To capitalize on the high temporal resolution of scalp EEG, we tested this independent mnemonic classifier on twenty 100 ms intervals across the List 2 stimulus interval (Figure 5). We find greater retrieval state evidence for OAs compared to MAs on encode trials and greater retrieval state evidence for MAs compared to OAs on retrieve trials. We tested these dissociations in a 2*×*2*×*20 mixed effects ANOVA with factors of age (MA, OA), instruction type (encode, retrieve) and time interval. We report the results of this ANOVA in Table 3; we find a significant three-way interaction between age, instruction type, and time interval, indicating that MAs and OAs differentially engage mnemonic brain states over time.

**Figure 5.**
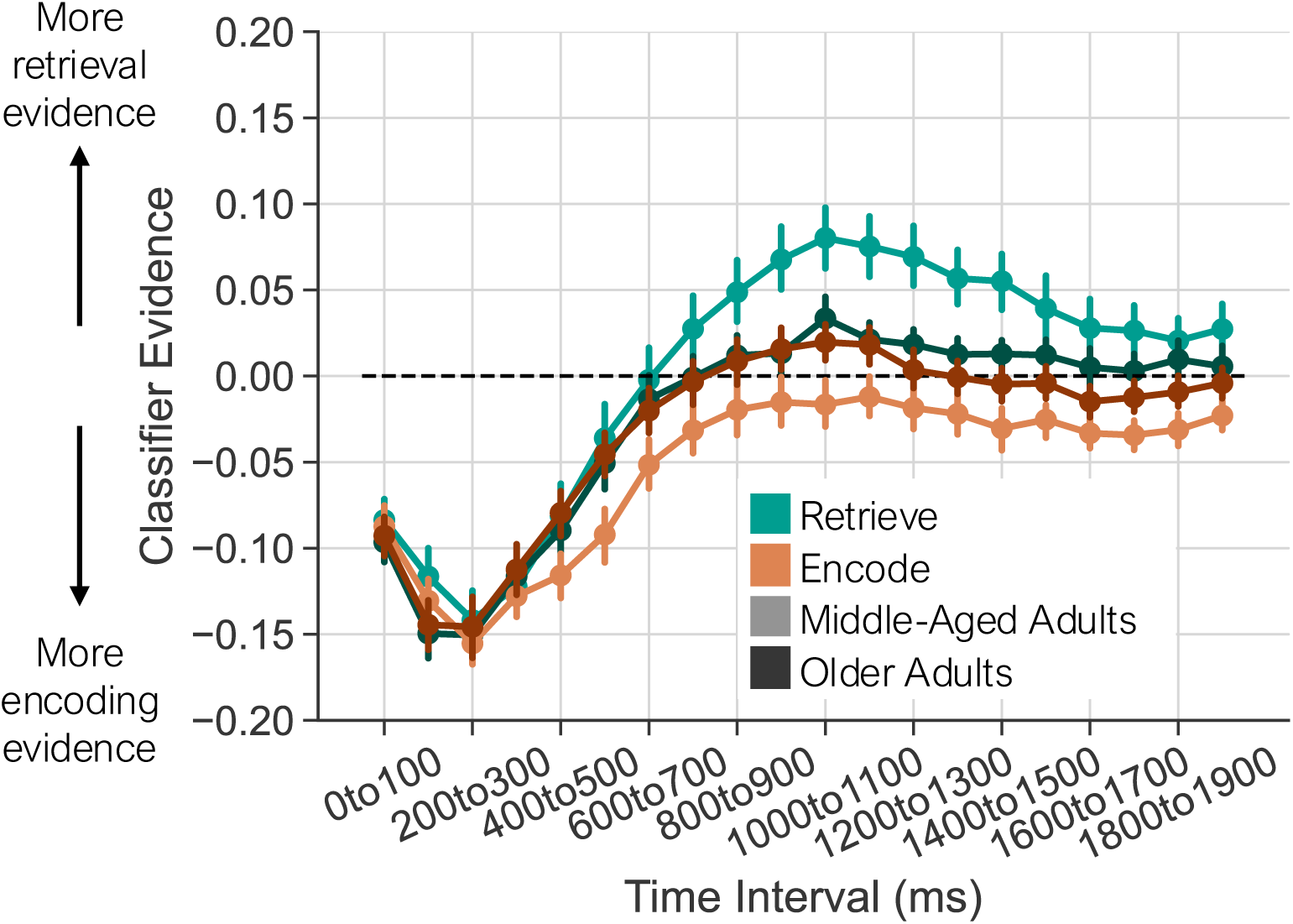
Independent assessment of mnemonic state engagement in middle-aged and older adults. We uti-lized an independently validated mnemonic state classifier to assess memory state evidence in twenty 100 ms intervals across the 2000 ms stimulus interval for both the middle-aged (lighter) and older (darker) adult data separately for encode (orange) and retrieve (teal) trials. We find a significant main effect of instruction type for middle-aged adults (*p <* 0.001), but not for older adults (*p* = 0.413). Error bars denote standard error of the mean.

**Table 3.**
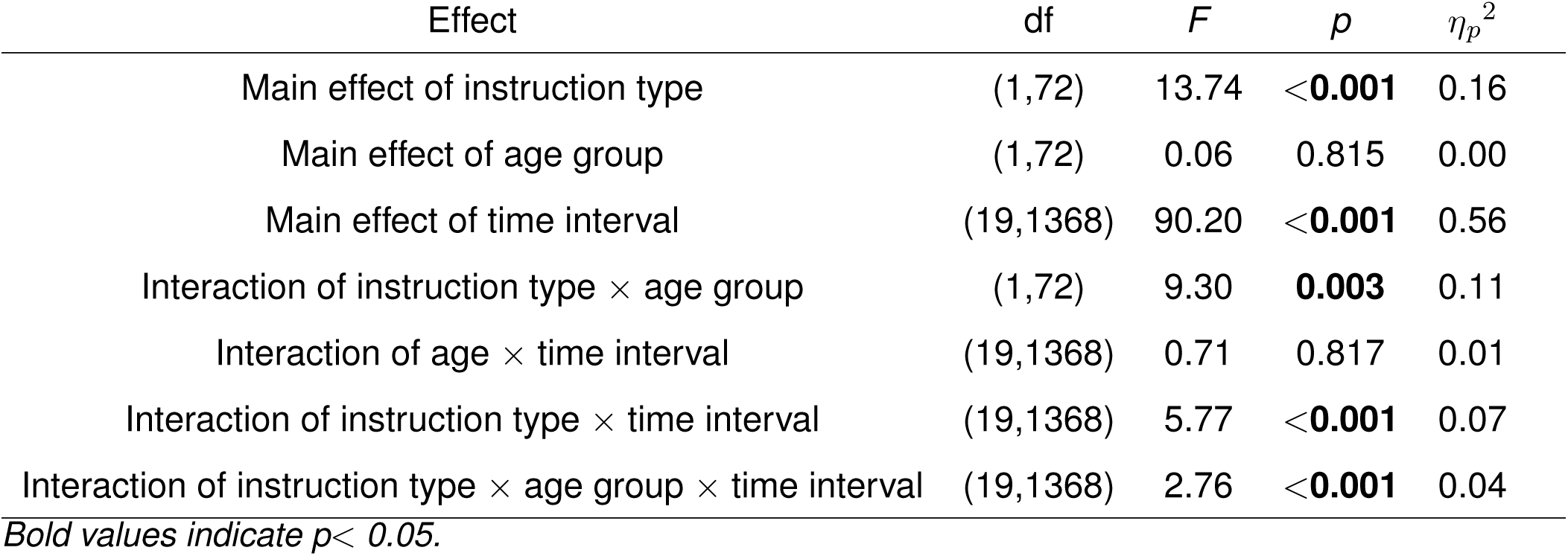
Analysis of variance results for instruction type, age group (middle-aged, older), and time interval (100 ms) on cross-study retrieval evidence.

| Effect | df | *F* | *p* | $\eta_p^2$ |
| --- | --- | --- | --- | --- |
| Main effect of instruction type | (1,72) | 13.74 | <**0.001** | 0.16 |
| Main effect of age group | (1,72) | 0.06 | 0.815 | 0.00 |
| Main effect of time interval | (19,1368) | 90.20 | <**0.001** | 0.56 |
| Interaction of instruction type × age group | (1,72) | 9.30 | **0.003** | 0.11 |
| Interaction of age × time interval | (19,1368) | 0.71 | 0.817 | 0.01 |
| Interaction of instruction type × time interval | (19,1368) | 5.77 | <**0.001** | 0.07 |
| Interaction of instruction type × age group × time interval | (19,1368) | 2.76 | <**0.001** | 0.04 |

*Bold values indicate $p < 0.05$ .*

We conducted two follow-up ANOVAs in which we compared mnemonic state engagement across instruction type and time interval separately for each age group. In MAs, we find significant main effects of time interval (*F* _19,665_ = 44.58, *p <* 0.001, *η_p_*^2^ = 0.56) and instruction type (*F* _1,35_ = 14.00, *p <* 0.001, *η_p_*^2^ = 0.29), the latter of which is driven by greater retrieval state evidence on retrieve compared to encode trials. We find a significant interaction between instruction type and time (*F* _19,665_ = 6.95, *p <* 0.001, *η_p_*^2^ = 0.17), whereby the greatest retrieval state engagement occurs during the middle of the stimulus interval on retrieve trials. In contrast, in OAs, we find a significant main effect of time only (*F* _19,703_ = 46.30, *p <* 0.001, *η_p_*^2^ = 0.56). We did not find a significant main effect of instruction (*F* _1,37_ = 0.69, *p* = 0.413, *η*^2^*_p_* = 0.02) nor a significant interaction between instruction and time (*F* _19,703_ = 1.03, *p* = 0.423, *η_p_*^2^ = 0.03). With the same training data, and hence, the same spectral features, these data suggest that mnemonic brain state engagement is diminished in OAs.

## Discussion

The goal of the current study was to measure the extent to which older adults engage in memory encoding and memory retrieval states. We directly tested the hypothesis that older adults are biased toward a retrieval state. We recorded scalp electroencephalography while young, middle-aged and older adult participants performed a memory task in which they were explicitly directed to either encode the currently presented object stimulus or retrieve a previously presented, categorically-related object stimulus. We report three key findings. First, we find that like young adults, middle-aged and older adults can follow top-down instructions to selectively engage in encoding or retrieval. Second, we find that we can successfully decode mnemonic brain states in all three age groups. Finally, we show that mnemonic brain state engagement is diminished in older adults as compared to middle-aged adults. Taken together, these results suggest that differential mnemonic brain state engagement for older adults compared to young and middle-aged adults may underlie age-related changes to memory.

Middle-aged and older adults selectively encode and retrieve in response to task demands. We find that across all age groups, memory is low for currently presented objects paired with a top-down demand to retrieve a previously presented object. As older adults have deficits in executive function (Foster, Cornwell, Kisley, & Davis, 2007; Plancher, Guyard, Nicolas, & Piolino, 2009; Salthouse, 2010, 2012), including difficulty inhibiting retrieval of semantic information (Wynn et al., 2020) and increased susceptibility to interference (Hasher & Zacks, 1988), we might have expected older adults to have worse memory performance for objects paired with the encode instruction relative to young and middle-aged adults. However, we did not find strong evidence for behavioral differences across age groups. As we did not directly assess retrieval success of the previously presented objects – by design, to minimize differences in behavioral demands between encode and retrieve trials – we cannot know what, if anything, participants retrieve during either encode or retrieve trials. Yet, given older adults’ tendency to rely on gist-level information rather than verbatim or specific details (Koutstaal & Schacter, 1997; Koutstaal, Schacter, Galluccio, & Stofer, 1999), older adults may have retrieved gist-level information, such as the object category, on encode trials. We expect that in contrast to verbatim retrieval of the previous item, gist-level retrieval would not interfere as strongly with the encoding of the current object given that it is from the same object category. Furthermore, as memory can be facilitated when older adults are aware of changes during new learning (Garlitch & Wahlheim, 2020), the impacts of interference might have been reduced in the current study given that we explicitly instructed participants as to the change in stimuli between the two object sets. Although there may yet be mechanistic differences as a function of age, our findings suggest that all three age groups can follow task instructions to selectively encode or retrieve.

We find that mnemonic brain states are decodable within each age group, demonstrating that middle-aged and older adults can shift between encoding and retrieval states in response to task goals. Given our hypothesis that older adults are biased toward a retrieval state, we may have failed to decode mnemonic states specifically for older adults. That is, if older adults engaged in retrieval on both encode and retrieve trials, the classifier would be unable to distinguish between those trials and mean classification accuracy would be at chance. That we find reliably above chance decoding in older adults provides evidence against our hypothesis and instead suggests that older adults can engage the encoding state despite potential interference from previously presented, categorically-related objects. This finding, like the behavioral results, may be the result of what is (or is not) retrieved. Older adults may automatically retrieve gist-level information without retrieving the specific previously presented, categorically-related object. In this case, there would be no internal target representation upon which to direct attention and thus, it might be easier for older adults to engage in encoding the externally presented stimulus. Interestingly, whereas classification accuracy was comparable between young and older adults, accuracy was greater for middle-aged adults. The increase in classification accuracy for middle-aged adults may be the result of an enhanced ability to sustain attention relative to young adults (Halder, Manot, & Chowdhury, 2019), although other work suggests that sustained attention abilities are stable across the lifespan (Carriere, Cheyne, Solman, & Smilek, 2010). Alternatively, middle-aged adults’ increased classification accuracy may arise from a combination of increased motivation and decreased sensitivity to stereotype threat. Specifically, although older adults may be more highly motivated than young adults when performing lab-based cognitive tasks (Frank, Nara, Touron, & Kane, 2015; Jackson & Balota, 2012; Seli et al., 2021), they may also be sensitive to stereotype threat – performance-related anxiety resulting from negative stereotypes about group mem-bership – resulting in poor task performance (Ryan & Campbell, 2021). Thus, middle-aged adults may benefit from the increased motivation associated with increased age, without the detrimental effects of stereotype threat found in older adults. Further work is necessary to disentangle the effects of sustained attention and motivation on classifier accuracy across age groups.

Despite finding above chance mnemonic state decoding in middle-aged and older adults, we find that mnemonic state engagement differs across age groups. Specifically, we find that compared to middle-aged adults, older adults show diminished engagement of both encoding and retrieval brain states. When we use an independent mnemonic state classifier to assess brain state engagement in middle-aged and older adults, we find two key dissociations between middle-aged and older adults. First, we find that older adults show diminished retrieval state engagement even when instructed to retrieve. Second, we find that older adults engage the retrieval state to a greater degree than middle-aged adults when instructed to encode. This latter finding is consistent with our hypothesis that older adults are biased toward retrieval. Our interpretation is that older adults have diminished executive control of brain state engagement as well as a modest bias toward the retrieval state.

Older adults may engage the retrieval state due to both a diminished ability to use top-down control to engage task-relevant mnemonic states and because automatic processes take over when control fails. Executive function deficits in healthy aging (Foster et al., 2007; Plancher et al., 2009; Salthouse, 2010, 2012) may result in a tendency toward the retrieval state. Specifically, researchers have posited that age-related memory declines may be the result of diminished executive functions with age (*executive decline hypothesis*; Dempster, 1992; Moscovitch & Winocur, 1992; Parkin & Walter, 1992; Troyer, Graves, & Cullum, 1994). The executive decline hypothesis is supported by evidence that older adults generally have worse performance on executive function tasks than young adults (Daigneault & Braun, 1993; Brennan, Welsh, & Fisher, 1997; Bryan, Luszcz, & Pointer, 1999) and is consistent with findings that older adults have difficulty with strategic encoding (Greene, Naveh-Benjamin, & Cowan, 2020) and inhibition (Hasher & Zacks, 1988; Connelly, Hasher, & Zacks, 1991; Hartman & Hasher, 1991; Hamm & Hasher, 1992; Hasher, Quig, & May, 1997; Lustig, Hasher, & Zacks, 2007; Lalla et al., 2022). Despite evidence that older adults appear able to engage mnemonic brain states to some extent, their engagement in these states appears diminished relative to middle-aged adults, which may be the result of age-related executive control deficits. These potential executive function deficits may be exacerbated by a baseline “shift” in the tendency to engage in semantic retrieval. Young adults automatically engage in retrieval when experiences overlap in time (Smith et al., 2022) and older adults have difficulty inhibiting retrieval from semantic memory (Wynn et al., 2020). In the event that older adults cannot sufficiently engage in either brain state, they might fall back on automatically engaged brain states. Given that older adults have greater DMN activity (Lustig et al., 2003; Grady, Springer, Hongwanishkul, McIntosh, & Winocur, 2006) and substantial prior knowledge, they may be automatically drawn into retrieval when they are unable to sufficiently engage encoding. Thus, older adults need to exercise a high degree of control – both to instantiate the task-relevant state and inhibit the task-irrelevant state.

Our interpretation that older adults’ executive control deficits may manifest as diminished mnemonic state engagement is consistent with a growing body of work suggesting that the mnemonic states measured here may reflect how attention is oriented rather than memory per se. That is, the current results may reflect attentional rather than – or in addition to – memory deficits. Attention can be divided into internal vs. external (Chun, Golomb, & Turk-Browne, 2011), whereby internal attention constitutes focusing on thoughts and representations in the mind whereas external attention constitutes focusing on sensory stimuli outside of the mind. Prior work has shown that the retrieval state as measured here is engaged in response to the top-down demand to retrieve (Smith & Long, 2024) in addition to spatial attention demands (Long, 2023) and changes in task demands (Wheelock & Long, 2024). Thus, older adults’ diminished engagement in memory encoding and retrieval states may reflect a decreased ability to deploy sustained attention to specifically focus their attention internally or externally.

In conclusion, we show that older adults’ mnemonic state engagement is diminished relative to middle-aged adults. Diminished mnemonic brain state engagement may arise from executive control deficits in older adults and may in turn account for episodic memory deficits in healthy aging. To the extent that the mnemonic states measured here reflect how attention is oriented and/or sustained, (Long, 2023; Smith & Long, 2024; Wheelock & Long, 2024), the age-related differences in mnemonic state engagement that we observe in the present study have a strong potential to influence cognition more broadly. These results advance our understanding of how mnemonic brain states change over the lifespan.

## Conflict of Interest

The authors declare no competing financial interests.

## Submission declaration and verification

This work has not been published previously and it is not under consideration for publication elsewhere. This publication is approved by all authors and will not be published elsewhere.

## Data statement

The raw, deidentified data and the associated experimental and analysis codes used in this study will be made available via the Long Term Memory lab website (https://longtermmemorylab.com) upon publication.

## Consent statement

All participants provided informed consent.

## Acknowledgments

We thank Yuju Hong for assistance with data collection. This work was supported by the National Institute on Aging of the National Institutes of Health under Award Number F31AG081045 (to ILM) and the integrative Translational Health Research Institute of Virginia (iThriv) Scholars Program (to NML). The iTHRIV Scholars Program is supported in part by the National Center for Advancing Translational Sciences of the National Institutes of Health under Award Numbers UL1TR003015 and KL2TR003016.

## CRediT authorship contribution statement

**Isabelle L. Moore:** Conceptualization, Data Curation, Formal analysis, Funding acquisition, Investigation, Software, Validation, Visualization, Writing – original draft, Writing – review & editing. **Devyn E. Smith:** Writing – review & editing, Investigation, Methodology, Software, Validation, Visualization. **Nicole M. Long:** Conceptualization, Formal analysis, Funding acquisition, Methodology, Project administration, Resources, Software, Supervision, Validation, Visualization, Writing – review and editing.

## Notes

### Competing Interest Statement

The authors have declared no competing interest.

